# Vimentin molecular linkages with nesprin-3 enhance nuclear deformations by cell geometric constraints

**DOI:** 10.1101/2024.10.29.621001

**Authors:** Maxx Swoger, Minh Tri Ho Thanh, Fitzroy J. Byfield, Van Dam, Jessica Williamson, Bronson Frank, Heidi Hehnly, Daniel Conway, Alison E. Patteson

## Abstract

The nucleus is the organelle of the cell responsible for controlling protein expression, which has a direct effect on cellular biological functions. Here we show that the cytoskeletal protein vimentin plays an important role in increasing cell-generated forces transmitted to the cell nucleus, resulting in increased nuclear deformations in strongly polarized cells. Using micropatterned substrates to geometrically control cell shape in wild-type and vimentin-null cells, we show vimentin increases polarization and deformation of the cell nucleus. Loss of nesprin-3, which physically couples vimentin to the nuclear envelope, phenotypically copies the loss of vimentin, suggesting vimentin transmits forces to the cell nucleus through direct molecular linkages. Use of a fluorescence resonance energy transfer (FRET) sensor that binds to the nuclear envelope through lamin-A/C suggests vimentin increases the tension on the nuclear envelope. Our results indicate that nuclear shape and deformation can be modified by the vimentin cytoskeleton and its specific crosslinks to the cell nucleus.

## Introduction

The architecture of the cell nucleus regulates nearly every facet of the cell, yet many aspects of how cells tailor nuclear shape remain unclear. While the nucleus has a smooth regular appearance in most resting cell types, cell migration in tissues involves polarization and large deformations of the cell nucleus to squeeze through small spaces and past neighboring cells [1–5]. The mechanical stresses associated with three-dimensional 3D cell migration can be sufficiently large to induce nuclear damage, in the form of DNA breaks, nuclear envelope rupture, and nuclear blebs, herniations of the nuclear envelope [6, 7]. Many pathologies are associated with abnormal nuclear shapes, which are commonly considered to arise from alterations in the mechanical properties of the nucleus itself or its ambient cell and tissue. For example, mutations in the nuclear lamins give rise to laminopathies, characterized by softer nuclei with many surface wrinkles and blebs [8–10]. Further, cancerous tissues are often abnormally stiff compared to healthy tissue and are also associated with nuclear blebbing and increased genome variation that stem from increased DNA damage [11].

Here, we examine whether the vimentin cytoskeleton is essential to the mechanisms by which cells control nuclear shape. In our prior study, we identified that vimentin protects the cell nucleus from damage during 3D cell migration through confining spaces and collagen networks [7]. Vimentin is a type III intermediate filament protein that forms a perinuclear cage, though its role in defining nuclear shape is not yet clear. Vimentin is also commonly used as a marker of the epithelial-to-mesenchymal transition, in which nonmigratory epithelial cells lose cell-cell adhesions and transition to a highly migratory mesenchymal phenotype [12]. Further, vimentin is expressed in many different cancer types and its expression level correlates with aggressive tumor growth and poor patient prognosis [13, 14]. These and other data suggest vimentin serves as a physical cushion that safeguards against large nuclear deformations during migration [7, 15]. This picture is further supported by the mechanical properties of purified and reconstituted vimentin networks, which are soft at small strains but significantly stiffen and resist deformations at large strains, giving the composite cytoskeleton optimal properties that cannot be achieved by F-actin and microtubules alone.

In this manuscript, we show that – in contrast to some earlier studies – vimentin enhances nuclear deformation and polarization, when cells are highly elongated and in a mechanically stressed state. Here, we focus on the effects of vimentin on nuclear shape when cell shape is geometrically constrained by micropatterned substrates. The cell geometric constraints allow control of cell shape, which regulates nuclear shape [16], cell contractility [17], and actin organization [18]. Our results highlight that the vimentin cytoskeleton plays an important role not only in mediating nuclear deformations induced by external or applied forces, but also aids in the transmission of the internal actomyosin networks that elongate the cell nucleus, a central feature behind the polarization and migration of cells. Since nuclear shape and positioning are essential to diverse functions in the cell, understanding the role of vimentin is important for interpreting many cellular mechanisms behind cell polarization, cell motility, wound healing, and cancer invasion.

## Results

### Vimentin enhances nuclear deformation on micropatterned substrates

To determine the effects of vimentin on nuclear shape, we studied wild-type and vimentin-null mouse embryonic fibroblasts (mEF) on collagen I-coated micropatterns (Methods), which geometrically constrain cell shape. The vimentin-null mEF cell line model expresses similar levels of actin and microtubules compared to wild-type cells but has no other cytoplasmic intermediate filament proteins [19, 20]. We focus on two geometric shapes: a square and a high aspect ratio (1:10) rectangle (Fig 1) that have the same surface area (300 μm^2^) yet result in two very different nuclear shapes [21]. On the square patterns, the cell nuclei are largely round, whereas on the rectangular pattern, the nuclei are ellipsoid. Strikingly, we find a notable difference in ellipsoidal nuclear shape in the wild-type and vimentin-null cells, as seen in Fig. 2. Namely, vimentin increases the nuclear elongation on the rectangle patterns, with an aspect ratio of approximately 2.39 in wild-type cells and 2.02 in vimentin-null cells (p<0.001). On square patterns, cells with and without vimentin had similarly shaped nuclei, as determined from the 2D projected images.

**Fig. 1:**
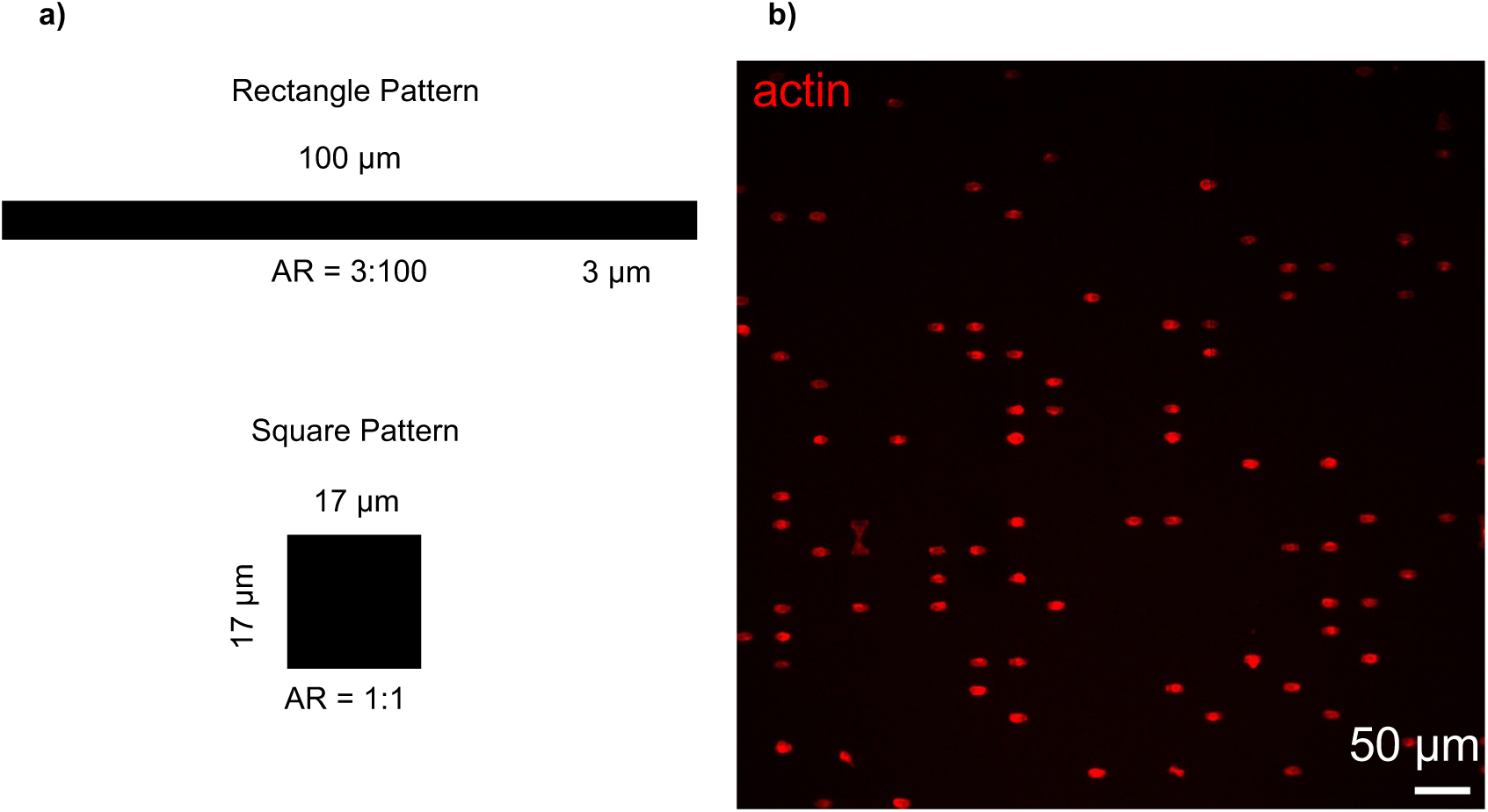
Micropatterns for cell geometric confinement. a) high aspect ratio micropattern 3:100 (rectangles) and low aspect ratio pattern 1:1 (squares) are used to geometrically confine cells. The square pattern serves as a control where we would not expect high strain on the nuclear envelope, whereas rectangles are expected to induce nuclear strain due to the high aspect ratio and width of 3 microns, less than half the radius of a typical nuclei. b) characteristic image of cells spread on a micro patterned glass coverslip; cells are stained with rhodamine phalloidin to visualize cell area. Image acquired with a 10x microscope objective.

**Fig. 2:**
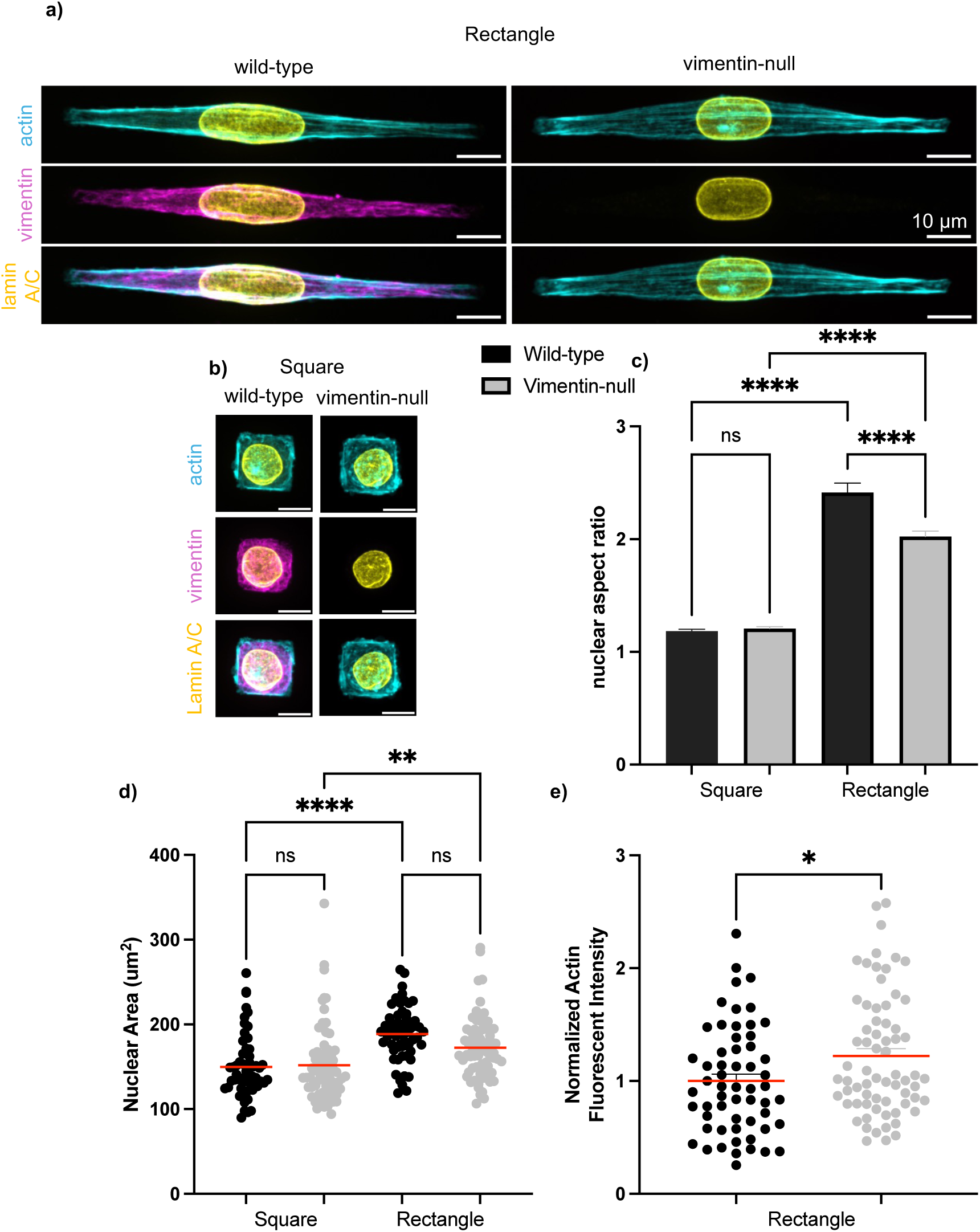
Vimentin enhances nuclear deformation on high aspect ratio patterns. Representative images of cells spread on rectangle a) and square b) patterns are provided where we show actin in cyan to show cell area, vimentin in magenta to show network structure, and lamin-A/C in yellow. Panel c) shows the nuclear aspect ratio found by tracing the lamin-A/C staining and panel d) shows the nuclear area. Actin fluorescent intensity was measured by tracing the cell area, the reported value is the mean fluorescent intensity where intensity has been normalized to wild-type cells for each experiment. All data represents N=2 experiments with n >= 20+ cells measured per condition. The statistical test is a two-way ANOVA with Šídák’s multiple comparison test for c) and d) and the data in e) was tested with an unpaired t-test with Welch’s correction.

Similar increases in nuclear deformation with intermediate filaments have previously been reported [22, 23]. Vimentin significantly increases the stiffness of cells under large mechanical loads, as shown by atomic force microscopy [24], suggesting vimentin is part of the molecular linkages that transmit forces to the nucleus. On the other hand, vimentin resists nuclear deformations for cells migrating through collagen networks [7]. Further, vimentin-mediated differences in nuclear shape have recently been argued to stem from differences in cell shape and were predicted to decrease nuclear deformation for the same cell shape based on the suggestion that vimentin protects the nucleus from large strains, the opposite of what we find here [25]. Taken together, there is conflicting data on whether vimentin enhances or hinders nuclear deformation and under what conditions. Here, we will present data that vimentin plays a significant role in increasing nuclear deformation in polarized cell states, namely through molecular linkages with nesprin-3 to the nuclear envelope.

First, however, we considered whether vimentin-mediated changes in cell or nuclear mechanics could account for the differences in nuclear shape, *e.g*. differences in actomyosin activity or nuclear stiffness (Fig. 2e). As cells initially make contact with a surface and spread, actomyosin networks generate contractile forces that allow cells to move and change shape. The microtubule network balances actomyosin contractile forces by resisting compression: disruption of the microtubule network induces larger cell contractile forces that would lead to larger nuclear deformations [25]. Here, no large differences in microtubules (SI Fig 1) are observed in the immunofluorescence images of the wild-type and vimentin-null cells. Specifically, the F-actin mean intensity is higher for vimentin-null cells on high aspect ratio patterns. Quantification of the length of actin stress fibers on the rectangle patterns shows no mean difference in stress fibers (data not shown), the higher actin intensity found for vimentin-null cells on high AR patterns clashes with the lower degree of nuclear deformation we observe. This data suggests that differences in actomyosin contractility is likely not responsible for the enhanced deformation we observe. Further, we previously showed that for rigid substrates, like the ones used here, vimentin-null cells generate larger traction forces [25], consistent with studies showing that loss of vimentin increases actomyosin contractility through RhoA and GEF-1 [26], consistent with our measurements of mean actin fluorescent intensity. One limitation here is that we did not measure actomyosin contractility via traction force microscopy on square and rectangular patterns directly. Nonetheless, the currently available data predicts a greater nuclear deformation in vimentin-nulls based on increased actomyosin contraction. Our results in Fig. 2c are, thus, in many ways unexpected.

A softer nucleus could give rise to larger deformations under the same applied forces. The nuclear lamins are a central factor in nuclear deformability, and we have previously shown that wild-type and vimentin-null cells have similar levels of lamin A, B1/2, and C by quantitative immunoblotting [7]. We have further experimentally tested whether we could detect any differences in nuclear stiffness by using atomic force microscopy techniques on isolated nuclei from wild-type and vimentin-null cells, but no difference in nuclear stiffness was detected. Further, on the rectangular patterns, there was no difference in immunofluorescence intensity of lamin A/C between the two cell types (SI Fig 2). Taken together, there is no indication that vimentin softens the cell nucleus to explain the increased deformations.

To analyze the 3D structure of the cell and the nucleus, we used confocal stacks to generate 3D renderings of the cells on micropatterned substrates. Nuclear lamins were used to demark the 3D boundary of the nuclear envelope (Methods). We observe two large characteristic stress fibers that run parallel with the long edges of the rectangular pattern below the nucleus and the expected cap of actin above the nucleus. The actin above and below the nucleus likely applies pressure to the nuclear envelope as the actomyosin network contracts on the pattern. The shape correlating to the pattern leads to the transition from a rounded dome morphology we see on the square patterns to an elongated cylindrical nuclear shape characteristic of high AR patterns. Furthermore, we have identified nuclear bulges and wrinkles associated with actin and/or vimentin fibers, on top of the nucleus. The average nuclear volume of the wild-type mEF was approximately 18% greater than the vimentin-null mEF (p = 0.0009, Fig. 3b) for cells on the square shapes, consistent with prior measurements on unpatterned glass that showed larger nuclear volumes in cells with vimentin [7]. While no change in nuclear volume is detected for the wild-type mEF on the rectangle vs. square patterns, the vimentin-null cells, in contrast, show a significant increase in nuclear volume of approximately 15% (p = 0.0052, Fig. 3b). The increase in mean nuclear volume for the vimentin-null cell is sufficiently large to match the nuclear volume of the wild-type cells on patterns. This result suggests vimentin mediates nuclear volume in a cell-shape dependent manner and may help maintain nuclear volume against changes in nuclear shape.

**Fig. 3:**
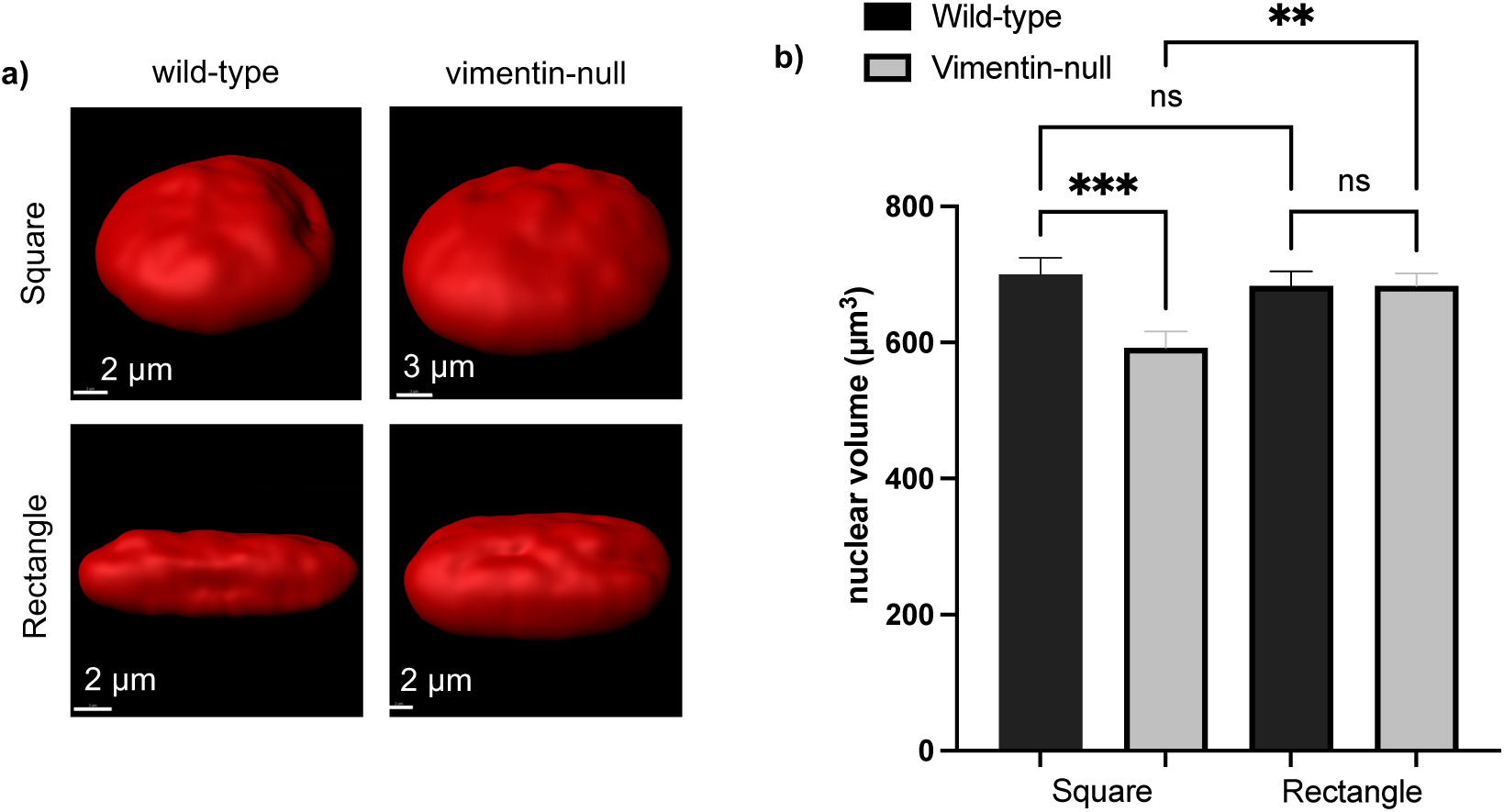
Determining nuclear volume from a 3D surface render of the nucleus. Panel a) shows characteristic surface renders of wild-type and vimentin-null mouse embryonic fibroblasts spread on either square or rectangular micropatterns. Surface renders are generated from confocal data where the entire cell nuclei has been imaged in 0.2 micron confocal slices, image processing and surface rendering was done with the software Imaris. b) shows the nuclear volume derived from the surface renders generated in Imaris. Data represents N = 2 experiments with n>= 20 cells per condition, statistical test is a two-way ANOVA with Šídák’s multiple comparison test.

### Molecular linkages between vimentin and the nuclear envelope increase nuclear deformations

Knocking out nesprin-3 in cells expressing vimentin results in reduced nuclear deformations, like cells with vimentin knocked out. Due to prior results from our group and others, our results are not fully explained by differences in actomyosin contractility or differences in nuclear stiffness between wild-type and vimentin-null mouse embryonic fibroblasts. Therefore, we chose to study the molecular links between vimentin and the nuclear envelope. The LINC complex is composed of a series of proteins that form a mechanical link that runs from the cytoskeleton to the chromatin, forming a mechanical link from which forces are transmitted into and out of the nucleus. Nesprin-3 is the linker protein associated with intermediate filaments that bond to the SUN 1/2 protein on the nuclear envelope. To investigate the role of the link between vimentin and the nuclear envelope we repeat our experiments on micropatterns with NIH 3T3 fibroblasts with and without nesprin-3, which both express vimentin.

Both 3T3 and N3KO cells were allowed to spread on square and rectangle patterns for 18 hours before fixing with 2% PFA, and nuclear and cell shape were visualized with an antibody against lamin A/C and rhodamine phalloidin, respectively (Fig. 4a and Fig 4b). We see a significantly higher nuclear aspect ratio for cells expressing nesprin-3 on high aspect ratio patterns, indicating that when nesprin-3 is deleted, vimentin’s effect on nuclear deformation is diminished (Fig 4c). From these results, we determine that nuclear deformations are enhanced by cells expressing vimentin only when vimentin is mechanically linked to the nucleus through the linker protein nesprin-3. The mechanical tether likely increased force transmission through vimentin to the nuclear envelope, resulting in larger deformations observed in our experiments.

**Fig. 4:**
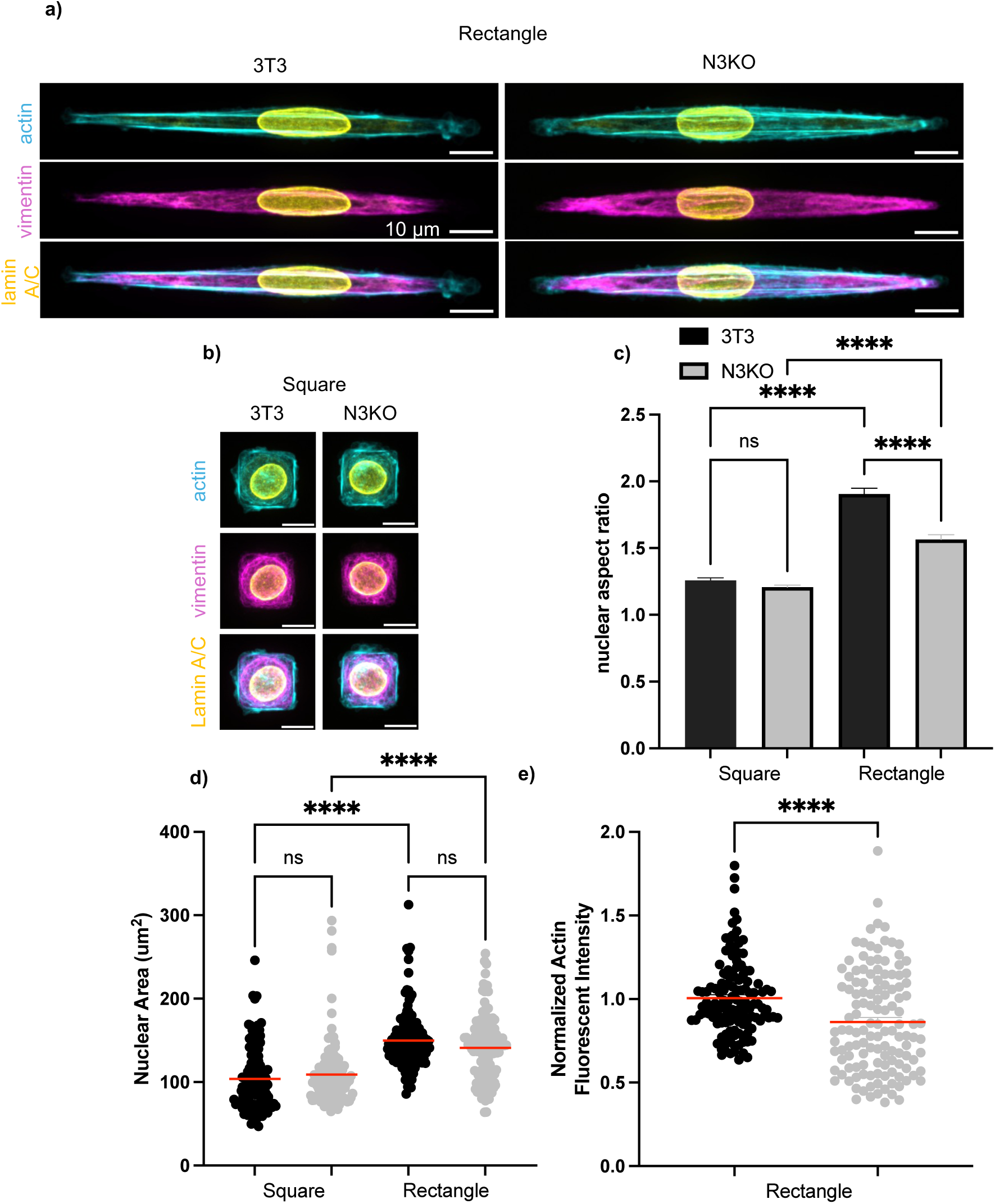
Nesprin-3 is required for enhanced deformation on high aspect ratio patterns. Representative images of 3T3 fibroblasts and 3T3 nesprin-3 knock out fibroblasts (N3KO) spread on rectangle a) and square b) patterns are provided where we show actin in cyan to show cell area, vimentin in magenta to show network structure, and lamin-A/C in yellow. Panel c) shows the nuclear aspect ratio found by tracing the lamin– A/C staining and panel d) shows the nuclear area. Actin fluorescent intensity was measured by tracing the cell area, the reported value is the mean fluorescent intensity where intensity has been normalized to wild-type cells for each experiment. All data represents N=2 experiments with n >= 20+ cells measured per condition. The statistical test is a two-way ANOVA with Šídák’s multiple comparison test for c) and d) and the data in e) was tested with an unpaired t-test with Welch’s correction.

Again, we sought to characterize how nesprin-3 knockdown could impact actomyosin contractility and nuclear stiffness. To assess the actomyosin contractility of the cells, we performed traction force microscopy on the wild-type 3T3 and nesprin-3 knockout cells (Methods). as shown in Fig. 5. We found that loss of nesprin-3 did not alter the mean traction force, suggesting similar levels of actomyosin contractility. Nuclear stiffness was measured by atomic force microscopy measurement of isolated nuclei. Likewise, no change in the mean nuclear stiffness was detected between the wild-type 3T3 and nesprin-3 knockout cells (Fig. 6). Taken together, these data suggest that nesprin-3 is important for nuclear elongation in polarized cells and this effect occurs even in the absence of notable changes in overall cell actomyosin contractility and nuclear stiffness.

**Fig. 5:**
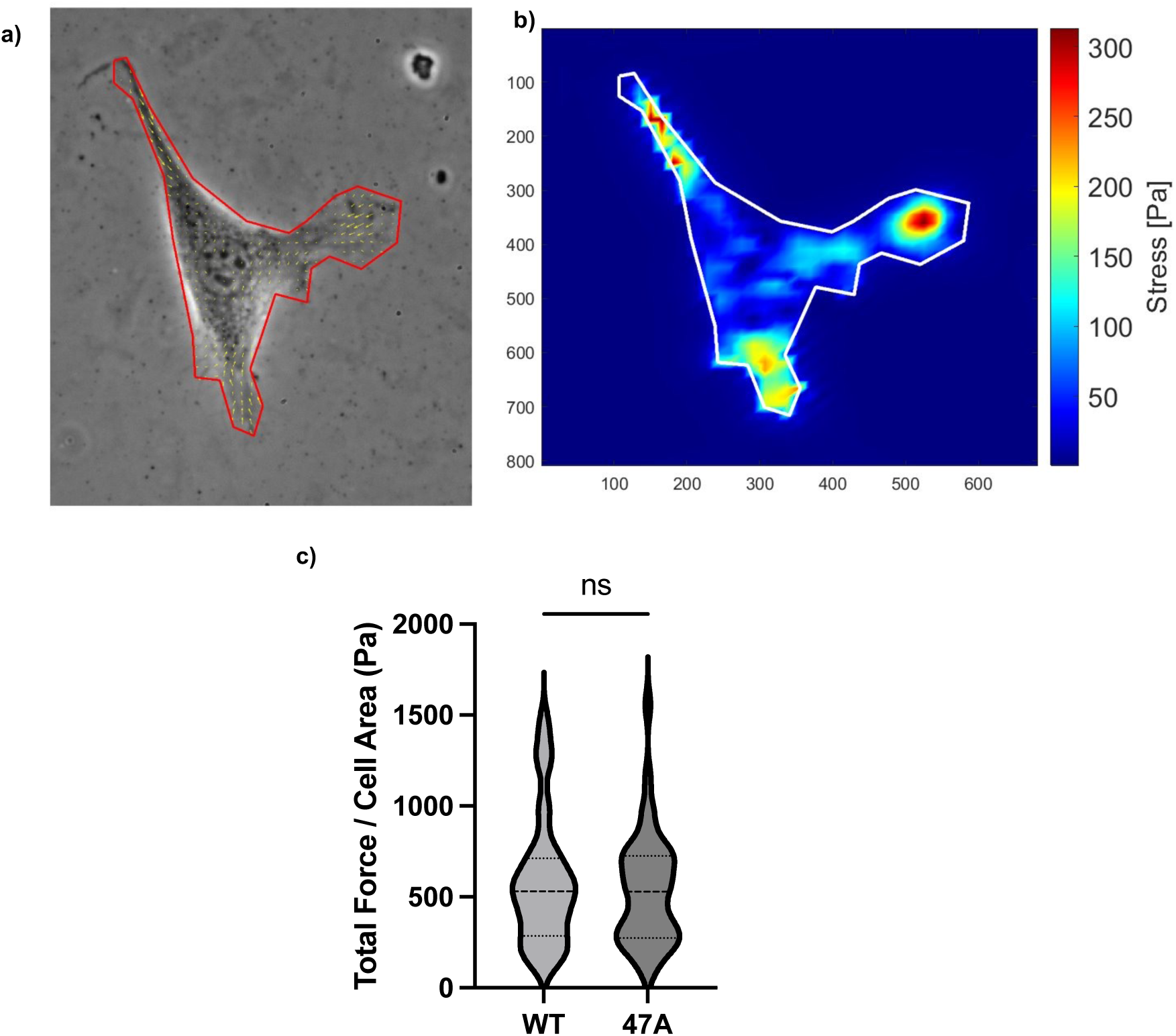
Characterization data for nesprin-3 knockout fibroblasts (N3KO) a) and b) show a nesprin-3 knockout fibroblast bead displacement field and traction stress map, respectively. c). Shows traction force microscopy data for 3T3 and N3KO cells, traction force measurements were performed on an 8 kPa polyacrylamide gel. (N=3 experiments, n=40+ cells total). Statistical significance was determined using a rank sum test.

**Fig. 6:**
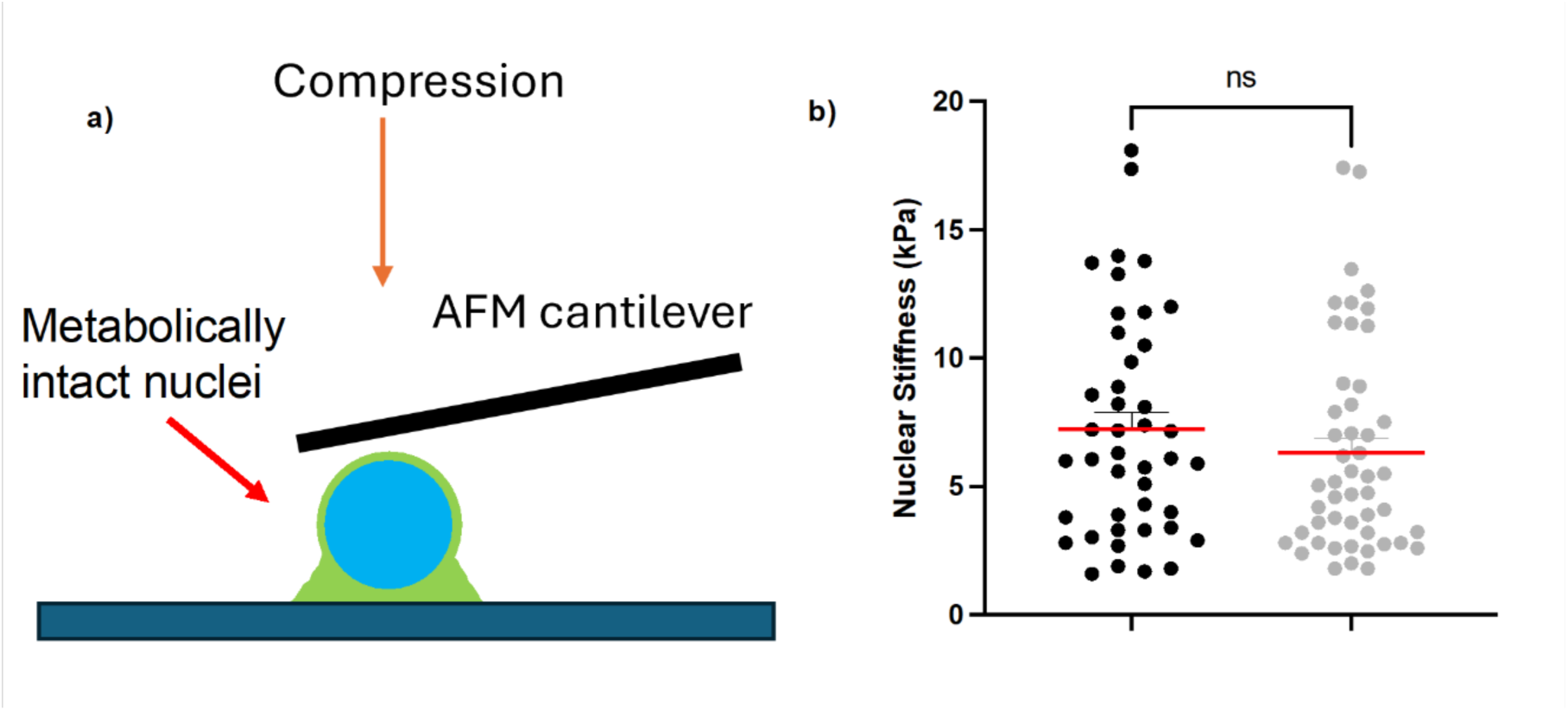
Characterization data for isolated nuclei of nesprin-3 knockout fibroblasts (N3KO. a) is a schematic representation for experiments that probe the isolated, but metabolically intact, nuclei of nesprin-3 knockout fibroblasts and 3T3 cells using a tipless AFM cantilever. b) shows the stiffness of nuclei that were isolated from either 3T3 or N3KO fibroblasts. Isolated nuclei are obtained by centrifugation, deposited on a PDL coated coverslip, and allowed to attach. These nuclei remain metabolically active after isolation. (N=3 experiments with n=60+ total nuclei per condition). Statistical significance for a) was tested with an unpaired t-test with Welch’s correction.

### Vimentin mediates chromatin condensation from nuclear elongation

Motivated by our findings that vimentin mediates changes in nuclear shape and volume, we hypothesized that disrupting vimentin may lead to alterations in chromatin configuration. To analyze chromatin configuration, we analyzed immunofluorescence images of wild-type and vimentin-null cells labeled with antibodies against trimethylation of lysine 9 on histone H3 (H3K9me3), a constitutive marker of heterochromatin (Fig. 7). Fig. 7a shows representative images of H3K9me3 on square and rectangle patterns for both cell types. The images show the characteristic foci of H3K9me3 domains that demark regions of heterochromatin in the nuclei. To quantify the H3K9me3 images, we measured the mean immunofluorescence intensity from cell nuclei (Fig. 7b) and the mean immunofluorescence intensity from individual foci in the nuclei (Fig. 7c). We found that the mean H3K9me3 nuclear intensities was greater for wild-type cells than vimentin-nulls on both the square and rectangle patterns, though the effect was stronger on the polarized rectangle shaped compared to the square pattern. The same pattern of results was found when we measured the intensity of individual foci. Interestingly, the H3K9me3 nuclear and foci intensities did not significantly vary for the wild-type cells on square vs. rectangles, but the vimentin-null mEF data showed a loss of H3K9me3 signal on the rectangle pattern compared to the squares (p < 0.0001). These results indicate that vimentin increases the level of heterochromatin in the cell nuclei and vimentin disruption leads to a loss of heterochromatin in a polarized state compared to an unstressed square shape, whereas wild-type cells maintain levels of heterochromatin upon changes in the nuclear shape observed here.

**Fig. 7:**
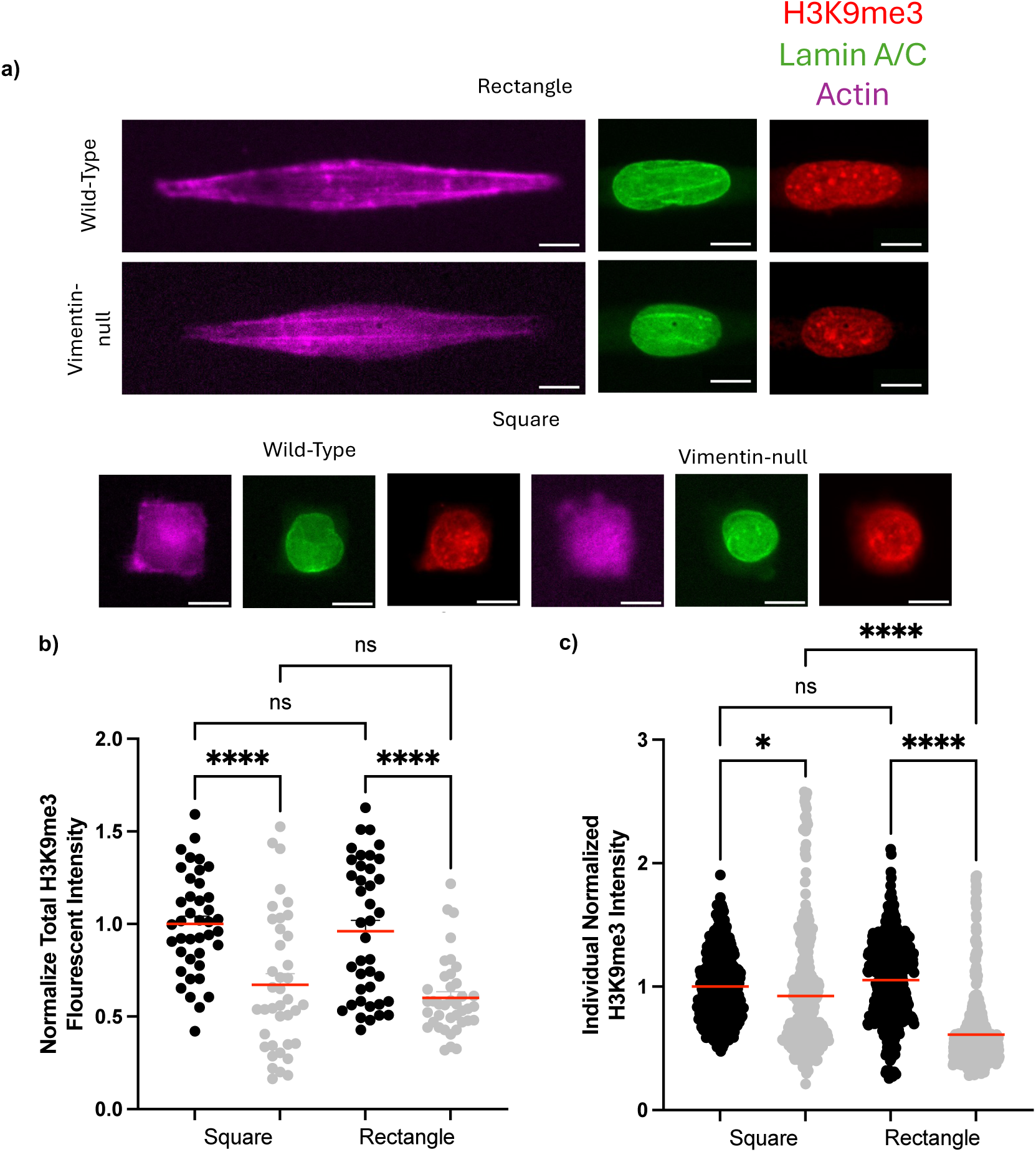
Expression of vimentin increases methylated DNA. a) shows staining of wild-type and vimentin-null fibroblasts spread on square and rectangular micropatterns where H3K9me3, a marker for methylated DNA is shown in red, lamin-A/C is shown in green, and actin is shown in magenta. b) shows the normalized total H3K9me3 fluorescent intensity found by tracing the nucleus and reporting the mean fluorescent intensity (N=2 experiments with n = 20+ cells). c) shows the mean fluorescent intensity of individual histones (N=2 experiments where all histones in 4 cells are traced n = 20+ histones per cell analyzed).

### Vimentin increases nuclear envelope tension

The experiments outlined so far in this work have focused on studying the shape of the nuclear envelope to make observations about the forces on the nuclear envelope. To directly probe the forces, we use a fluorescence energy transfer (FRET) sensor for lamin A/C (Fig 8a). This probe is a stretchable, mechanically sensitive, sensor with either end presenting chromobodies for lamin A/C, so that the sensor localizes to lamin A/C when expressed. Wild-type and vimentin-null fibroblasts were transfected with either a tension sensing or truncated mutant (control) version of the lamin A/C FRET Sensor (Fig 8b).

**Fig 8:**
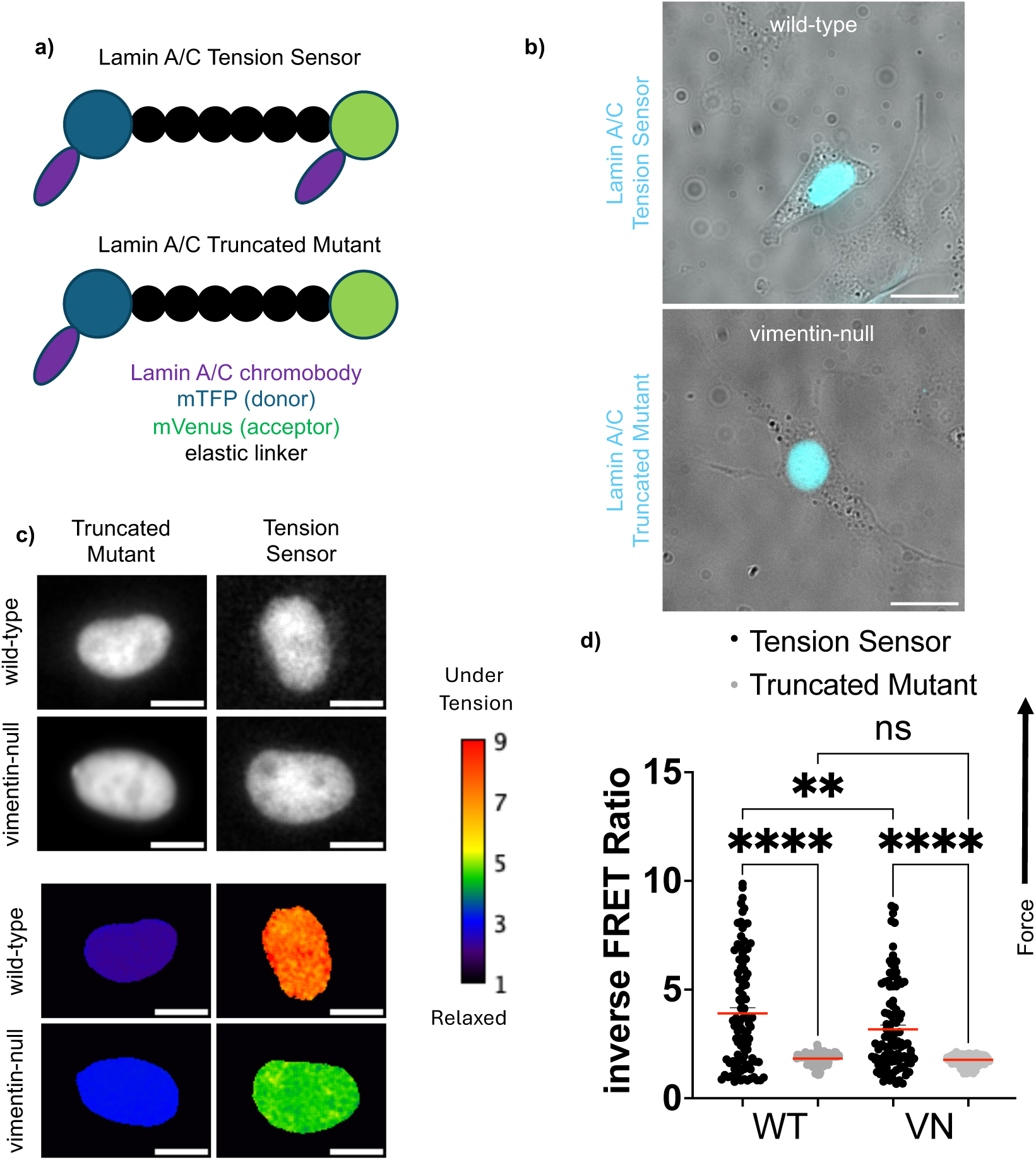
Cells expressing vimentin have higher relative stress on their nuclear envelope. (a) shows a schematic of the lamin-A/C fluorescent resonance energy transfer (FRET) sensor that is transfected into wild-type and vimentin-null fibroblasts allowed to freely spread on a glass coverslip (b). There are two types of sensors employed, a tension sensitive one which contains both lamin-A/C binding domains and a control tension insensitive truncated mutant where one of the lamin-A/C binding sites is cleaved off, resulting in the sensor being stuck in a relaxed or high FRET state. c) shows representative images of single fluorophore images (TFP channel) to show localization of the sensor to the nuclear envelope. c) are characteristics color maps of the inverse FRET ratio, where a hotter color represents a region under increased levels of strain. d) shows quantification of FRET data where we plot the inverse FRET ratio, meaning a higher value indicates higher relative forces or strain on the nuclear envelope. (N=2 experiments with n = 50+ histones per cell analyzed, outliers are removed by identifying values falling outside of the range two standard deviations from the mean). The statistical test is a two-way ANOVA with Šídák’s multiple comparison test.

Cells were allowed to spread on glass bottom culture dishes post-transfection for 18 hours before fixing with 2% PFA. Two sets of images were collected for each cell: a YFP channel image and a CYP->YFP bandpass image. Analysis of FRET data was performed by following the protocols for ratiometric FRET published previously [27, 28]. Images undergo a background subtraction, are converted to 32-bit, and smoothed. A threshold is generated using the YFP channel to mask out any data outside of the nuclear envelope. Inverse FRET ratio was calculated by dividing the CFP-> YFP image by the YFP image (Fig 8c). We checked sensor expression by seeing localization to the cell nuclei and comparison of the tensor sensing and truncated mutant sensors. We observe that wild-type cells have a significantly higher relative force on their nuclear envelope than vimentin-null cells (Fig 8d). This suggests that vimentin enhances force transmission to the nuclei, consistent with the higher degree of nuclear deformation we observe for wild-type mouse embryonic fibroblasts.

## Discussion

Nuclear structure is highly regulated within the cell, but the mechanisms that sculpt nuclear shape under different conditions remain unclear. Our results here show that vimentin is important for facilitating the forces that elongate the cell nucleus in geometrically constrained polarized cells. Further, depletion of nesprin-3 disrupts the elongation of the cell nucleus, suggesting vimentin’s physical linkages to the nuclear envelope are involved in this deformation of the cell nucleus.

Vimentin is traditionally perceived as a passive cytoskeletal network that does not interact with molecular motors, in contrast to the two other cytoskeletal polymers, F-actin and microtubules. One could reason that vimentin’s physical crosslinks with F-actin and microtubules and its impact on cell stiffness would enhance the transmission of cell-generated forces to deform the nucleus, as we find in Figure 2. However, the strong impact of nesprin-3 knockout in cells with vimentin suggests vimentin alone is not sufficient to deform the cell nucleus. Vimentin’s molecular linkages to the nuclear envelope are critical to elongating the nucleus in these polarized cells.

The ability of vimentin to increase cell-generated nuclear deformations may be surprising, given findings that vimentin plays an antagonistic role with actomyosin contractility [26, 29]. Previous studies by Jiu et al showed vimentin mediates actin stress fiber assembly through the microtubule-associated guanine nucleotide exchange factor GEF-H1 that activates RhoA, decreasing actomyosin contractility and cellular traction forces[26]. These studies suggest that loss of vimentin would increase actomyosin activity, leading to larger nuclear deformations, in contrast to what we find on the rectangular patterns used here. Recent works from our group and others have shown that vimentin’s effects on cellular traction stresses are dependent on stiffness of the underlying substrate [29, 30]. These studies show that vimentin decreases actomyosin contractility on stiff substrates (approximately > 20 kPa) but increases actomyosin contractility on soft substrates (< 20 kPa). This data is consistent with a chemomechanical model in which vimentin assists with actomyosin contractility by forming physical crosslinks with the network that increases the transmission of those forces on soft substrates. On the other hand, on stiff substrates, where cell-generated stresses are much larger, vimentin plays a larger role in supporting the microtubule network, which resists actomyosin contractility, by serving as a polymer network that buttresses and prevents breakage and buckling of microtubules. In this study, cells are on micropatterned glass substrates, which are effectively infinitely stiff to the cell, a regime in which it would be predicted that loss of vimentin increases nuclear deformation. Our results thus highlight an effect of vimentin on nuclear shape that is unexpected in many ways, and our results suggest this effect is dependent on physical crosslinks between vimentin and the nuclear envelope through nesprin-3. Together, our results suggest new models of nuclear deformation are needed that explicitly account for the effects of vimentin and its molecular linkages to the nuclear envelope.

Vimentin intermediate filaments are important for cell polarization [31, 32]. Vimentin presents a template for microtubule network structure persisting through cycles of disassembly and reassembly, aiding in maintaining cell polarization [32]. For unpatterned cells on 2D soft substrates, vimentin increases the orientation of traction stresses, enhancing the polarization of the cell [31]. Interestingly, recent studies show vimentin aids the transition from a collective cell jammed to unjammed state in cancer cells and fibroblasts, resulting in an increased polarized migratory phenotype [33]. These studies all point to an increase in cell polarization via vimentin, which could contribute to the processes that polarize and elongate the cell nucleus as seen here.

Our results suggest vimentin plays a complementary role with nuclear lamins in mediating nuclear shape. The nuclear lamins are important to preserving nuclear shape. Disrupting nuclear lamins increases nuclear deformations and can lead to nuclear abnormalities, such as blebbing and nuclear envelope rupture [9, 34–36]. Loss of specific lamin isoforms impacts cortical and cytoplasmic stiffness [37] as well as nuclear deformability, particularly at large nuclear strains [38]. Further, Vahabikashi et al [37] showed that nuclei in lamin knockout or knockdown MEF lines were significantly rounder than those in wild-type mEF. We note that we have previously shown that loss of vimentin does not change the expression levels of lamin A/C or lamin B [7] nor modulate nuclear stiffness, as measured by atomic force microscopy measurements of isolated nuclei [24]. Given the importance of maintaining the structural integrity of the nucleus, it stands to reason that the cell has multiple networks to fine-tune the shape and deformability of the nucleus.

Our data demonstrates a clear (and perhaps unexpected) link between vimentin and nuclear shape in polarized cells. The interactions between vimentin and the cell nucleus are clearly complex and likely a strong function of both mechanical and non-mechanical effects. It is tempting to want to simplify vimentin’s effects to a primary biological function, but as we see here, vimentin can serve many different functions under different conditions. This is in part because of the many phosphorylated states of vimentin that trigger very different downstream biological signals [39]. The diversity of structure and functions to allow the cells to adapt to different environmental conditions is thus a salient feature of intermediate filament proteins.

Our data here provides new evidence for the role of vimentin in nuclear volume and chromatin organization. Nuclear volume is known to increase with cell volume [40], but wild-type cells had larger nuclear volume than vimentin-null cells of the same cell volume (on unpatterned substrates). Likewise, we find that nuclear volume is greater for wild-type cells on the square patterns. Here, we show that cells with vimentin maintain nuclear volume despite changes in nuclear shape (square vs. rectangles), whereas vimentin-null cells show increased nuclear volumes on rectangle patterns. Vimentin may thus work to resist changes in nuclear volume based on varying cell shapes.

The data on H3K9me3 show that there is more heterochromatin in cells with vimentin. A prior study by Alisafaei, et al, indicates a connection between nuclear volume and heterochromatin levels, mediated by actomyosin contractility on different pattern types [41]. Specifically, larger and higher-aspect ratio patterns were correlated with higher levels of actomyosin markers, such as phosphorylated myosin light chain immunofluorescence. The higher actomyosin contractility led to increased nuclear-to-cytoplasmic translocation of histone deacetylase 3 (HDAC3), causing chromatin decondensation and increased nuclear volume. An interesting observation is that some of the predicted effects are observed here, *e.g.* increased nuclear volume on rectangles for vimentin-null cells, while others are not, *e.g.* heterochromatin levels do not strongly correlate with nuclear volume for either cell type. This data suggests there may be other unrecognized factors in mediating nuclear shape, and there is an interesting but not fully resolved link between vimentin and nuclear volume in our finding. The connection between vimentin and nuclear shape is also seen in recent studies [24] that show vimentin increases nuclear volume in whole cells, but in isolated nuclei, wild-type and vimentin-null cells have the same volume. These suggest there is a possible mechanism by which the vimentin perinuclear cage increases nuclear volume that may be independent of the nuclear envelope and structure itself.

Finally, our work highlights opposing results between vimentin protecting the nucleus during migration in 3D confining environments [6, 7], but we find larger deformation for cells on patterned 2D substrates for cells expressing vimentin mechanically linked to the nucleus. Interestingly, it was recently shown that when the cell nuclei undergo a confining strain it initiates an evasive response from the cell to escape the confinement [42]. Our studies show that vimentin enhances nuclear deformation in 2D, and we propose that vimentin may perform a similar function in 3D, leading to aversion to confining environments as vimentin triggers the mechanosensitive response of the nucleus, connecting the behavior we observe in 2D to our previous studies in 3D environments.

## Conclusion

In these experiments we observe nuclear deformations for wild-type and vimentin-null mouse embryonic fibroblasts, where we find that vimentin enhances nuclear deformation for cells on confining micropatterns. Furthermore, we demonstrate that vimentin must be mechanically linked to the nuclear envelope through the linker protein nesprin-3 for this deformation to be observed. To probe the relative forces exerted on the cell nuclei, we employed a FRET tension sensor that binds to Lamin A/C. The data indicated that wild-type fibroblasts have a higher relative force on their nuclei. Taken together, these results underscore the need to consider intermediate filaments, such as vimentin, in models for nuclear shape, chromatin configuration, and mechanosensing of the cell.

## Acknowledgements

We thank useful discussions from Jen Schwarz and Paul Janmey. This manuscript was supported by NSF CMMI 2238600 and NIH R35GM142963, awarded to AP, and NIH R35 GM119617, awarded to DC.

## Materials and Methods

### Cell culture

Wild-type and vimentin-null mEFs were kindly provided by J. Ericsson (Abo Akademi University, Turku, Finland). Cells were cultured in media composed of Dulbecco’s Modified Eagle’s Medium (DMEM) + 4.5 g/L glucose, L-glutamine, and sodium pyruvate (Corning). Culture medium was supplemented with 10% fetal bovine serum (Hyclone), 1% penicillin streptomycin (Fisher Scientific), 25 mM HEPES (Fisher Scientific), and 1% nonessential amino acids (Fisher Scientific). Cell cultures were maintained at 37 C with 5% CO_2_, cultures were passaged when they reached 70% confluency. NIH-3T3 cells were obtained from ATCC (CRL-1658).

NIH-3T3 nesprin-3 knockout cells were generated by using CRISPR/Cas9 by transfecting cells with an All-in-One U6-gRNA/CMV-Cas9-tGFP Vector (Sigma) using gRNA sequence: TTTGTAACCGGTCTACACG. Cells were transfected with the CRISPR/Cas9 plasmid (using Lipofectamine 2000) followed by clonal selection and expansion. Clonal isolates were screened for loss of nesprin-3 by Western blotting using an anti-nesprin-3 rabbit antibody (gift of Dan Starr, UC Davis). 3T3 cells were cultured in in media composed of Dulbecco’s Modified Eagle’s Medium (DMEM) + 4.5 g/L glucose, L-glutamine, and sodium pyruvate (Corning). Culture medium was supplemented with 10% bovine serum (Hyclone), 1% penicillin streptomycin (Fisher Scientific), 25 mM HEPES (Fisher Scientific), and 1% nonessential amino acids (Fisher Scientific). Cell cultures were maintained at 37 C with 5% CO_2_, cultures were passaged when they reached 70% confluency.

### Immunofluorescence

Cells were fixed for immunofluorescent staining using 2% paraformaldehyde (Fisher Scientic) for 30 minutes at 37 C. Cell membranes are permeabilized by exposure to 0.05% Triton-X (Fisher BioReagents) in PBS for 15 minutes at room temperature, then samples are blocked with 1% bovine serum albumin (BSA) (Fisher BioReagents) for 30 minutes at room temperature. Cells were incubated for 1.5 hours at room temperature with primary antibodies at a dilution of 1:200 in 1% BSA in PBS. For visualization of vimentin an anti-vimentin rabbit monoclonal antibody (Abcam) or anti-vimentin polyclonal chicken antibody. Lamin A/C was visualized using an anti-lamin A/C monoclonal mouse antibody (Cell Signaling). Samples were incubated with secondary antibodies for 1 hour at room temperature at a dilution of 1:1000 in 1% BSA in PBS. Donkey anti-rabbit Alexa Fluor 568 (Invitrogen), goat anti-rabbit Alexa Flour 488 (Invitrogen), or anti-chicken Alexa Fluor 488 (Invitrogen) were used for visualization of vimentin. Goat anti-mouse Alexa Fluor 647 (Invitrogen) or goat anti-mouse Alexa Fluor 488 (Invitrogen) were used for visualization of Lamin A/C. Cells were stained with Hoeschst 33342 (Molecular Probes) at a concentration of 1:1000 in 1% BSA in PBS for 1 hour to detect DNA. To visualize actin samples were stained with either rhodamine phalloidin 565 (Invitrogen) or rhodamine phalloidin 647 (Invitrogen) at a dilutions of 1:200 in 1% BSA in PBS for 1 hour. Samples were mounted using either ProLong Diamond Antifade (Life Technologies) or Vectashield Antifade Mounting Medium (Vector Laboratories) sealed with nail polish.

Cells were imaged using a Leica DMi8 (Leica) equipped with a spinning disk X-light V2 Confocal Unit using a 40x water immersion objective or 60x oil immersion objective, controlled with Visiview software (Visitron Systems). Images were acquired by collecting z-stacks with 0.2 micrometer step sizes, and data was analyzed with ImageJ (NIH).

### Substrate Preparation and cell plating

Uncoated micropatterned glass coverslips were obtained with the Humen-3 pattern from CYTOO (CYTOO). The CYTOO chip was coated with 40 ug/mL rat tail collagen I (Corning) for 2 hours at room temperature, then excess collagen is washed out by serial dilution with PBS. Collagen coated coverslips are stored in PBS at 4 C prior to experiment, at least 30 minutes prior to an experiment PBS is aspirated, replaced with cell media, and the dish containing the coated chips in put in the incubator at 37 C. Cells are disassociated from culture dishes by exposure to TrypLE Express Enzyme (Gibco) for 5 minutes, TrypLE is deactivated by addition of cell media when cells are suspended, cells are then transferred to a centrifuge tube and spun down for 2000 RPM for 5 minutes. The supernatant is aspirated, cells are resuspended, and then cells are counted using a Countess 3 Automated Cell Counter (ThermoFisher). A cell suspension with a concentration of 15,000 cells/mL is made and 4 mL of the cell suspension is added to each well containing a coated CYTOO chip. Cells are allowed to spread for 18 hours before fixation.

### FRET sensor expression

The FRET sensors used in these experiments were previously reported in Danielsson et al. [43]. There were two versions of the sensor employed, a force sensitive tension sensor and a control truncated mutant sensor, where both sensors facilitate binding to lamin-A through a nanobody, however the truncated mutant has one binding site deleted so that it is permanently in a relaxed state to serve as a control. The FRET pair used in these sensors is TSmod, where mTFP serves as the donor fluorophore and venus is the acceptor fluorophore.

Stocks of the plasmid for the truncated mutant and tension sensor plasmid were created using a Purelink HiPure Plasmid Filter Maxiprep Kit (Thermo Scientific) that exceeded a concentration of 1 μg/μL. Expression of sensors was induced in cells using the following procedure. Cells are allowed to grow to 80% confluence in a six well plate, wash cells with reduced serum media (Gibco), then place in incubator while preparing the transfection solutions. Prepare a lipomix by adding Lipofectamine 2000 (Invitrogen) to reduced serum media at a ratio of 1:40, then gently mix. The DNA transfection mix is prepared by adding 1 μg of DNA per well of an aliquot of reduced serum media, then adding an equal part lipomix and mixing gently. The transfection solution was incubated for 20 minutes at room temperature before 1 μg of DNA was transferred to each well.

Cells were incubated in transfection media for 18 hours, then washed twice with culture media, and expression of sensor was verified by examining on a fluorescent microscope using a GFP filter.

### FRET Image Acquisition and Analysi**s**

Prior to experiment, recovered cells are disassociated from culture dishes using TrypLE Express Enzyme (Gibco), which is then deactivated by adding cell media after cells are suspended. Cells are transferred to a centrifuge tube, spun down for 2000 RPM for 5 minutes, the supernatant is aspirated, cells are resuspended, and the cell density is found using an automated cell counter. Cells are plated on a 35 mm dish with a 28 mm glass coverslip bottom well (Cellvis) at a final count of 270,000 cells per dish. Cells are allowed to spread on the dish for 18 hours before being fixed with 2% PFA for 30 minutes at 37 C. The fixed samples are stored in PBS until imaging. Cells were imaged using a Leica DMi8 (Leica) equipped with a spinning disk X-light V2 Confocal Unit using a 40x dry objective and a pair of FRET imaging cubes, one to image just the YFP channel and the other to image the YFP->CFP FRET channel. Images were acquired using the VisiView software (Visitron Systems).

Analysis of FRET data was done by following protocols reported previously [27, 28], to generate ratiometric FRET images with the image analysis software ImageJ. The following steps are performed for both the YFP and YFP->CFP image. First a background subtraction using a rolling ball, then images are converted to 32 bit, and finally the images are smoothed. The YFP image is automatically thresholded and all values outside of the image threshold generated are changed to NaN so that they are ignored in the final FRET image map, meaning only FRET ratio values inside of the nucleus are recorded. Finally, an inverse FRET image map is generated by dividing the YFP->CFP image by the thresholded YFP image using the Ratio Plus plug in, and an average value is generated by using the measure feature and recording the mean.

### Imaris Analysis

Surface renderings of cell nuclei were generated with the surface volume feature of Imaris (Oxford Instruments). Confocal stacks of the entire cell volume were imaged with a 0.2 μm step size using a spinning disk confocal microscope as detailed above. Surface rendering of nuclei were generated using methods previously reported [7, 44]. In summary, images were opened in the Imaris software, and the image was cropped to a region slightly larger than the nucleus, and a surface was generated using the default parameters. The surface render uses a background subtraction for thresholding, where the input is the smallest diameter of the nuclei measured in the slice view of Imaris.

Finally, surfaces are inspected by eye for any artifacts and statistics about nuclear volume are exported from the Imaris software.

### Traction force microscopy

Polyacrylamide (PAA) gel embedded with fluorescent beads was made with the following protocol [45]. The surface of 25×25 mm glass coverslips (Corning) were cleaned and silanized with 1% 3-aminopropyl-trimethoxysilane solution (Acros Organics) for 10 min and then treated with 0.5% glutaraldehyde (Fisher Scientific). 18 mm round glass coverslips (Fisher Scientific) were plasma-cleaned and spin-coated with 0.25 μm fluorescent (515 nm/585 nm) beads (Spherotech) on the surface. Stock solutions of acrylamide and bis-acrylamide mixture were made by mixing an 200 μL of 40% acrylamide (Fisher Scientific) and 50 μL of 2% bis-acrylamide (Fisher Scientific) in 735 μL of HEPES buffer (50 mM pH = 8.2) to generate gels of Young’s moduli E= 8 kPa [45]. After degassing 985 μL of the mixture, 10 μL of 10% ammonium persulfate (Thermofisher) and 3 μL of N,N,N°,N°-tetramethyl ethylenediamine (Bio-Rad) were added to initiate the acrylamide polymerization. Immediately after the initiators were added, 25 μL of the solution was taken out and sandwiched between the bead-coated coverslip and the glutaraldehyde-coated coverslip. After 15 minutes of polymerization, the gel and coverslips system were then sub-merged in cold HEPES buffer for 30 min. The round coverslip was then carefully removed, resulting in a beads-coated gel disk of 18 mm in diameter and approximately 100 μm in thickness attached on the 25×25 mm square coverslip. Collagen type I was then crosslinked to the PAA gel surface using sulfo-SANPAH. The gels were submerged under 1 mg/mL sulfo-SANPAH (Thermofisher) solution and placed under UV for 10 minutes. The gels were then washed with PBS buffer and soaked with 0.1 mg/mL rat-tail collagen type I (Corning) overnight at 4°C. After collagen coating, the gels were rinsed with PBS buffer for 30 minutes. The collagen-coated gels were placed in culture medium and incubated for 15 min at 37°C before seeding cells on top.

Traction force microscopy (TFM) was performed on a Leica DMi8 using a 40x HC PL Fluotar (Leica) dry objective lens. Cells were cultured on PAA substrates for 12–16 hr before subjected to TFM. Only single cells were selected for TFM. For each cell, a phase contrast image of the cell and a fluorescent image of the beads were acquired using K5 sCMOS camera (Leica). Cells were then released from the substrate using 0.25% TrypLE (Thermofisher). After trypsinization, an image of fluorescent beads in the undeformed substrate is acquired again. The pair of fluorescence images were analyzed using a particle imaging velocimetry routine in MATLAB (MathWorks) to extract the displacement field on the PAA gel generated by cell traction force. Substrate material properties, substrate dimensions, bead displacements and cell boundary was imported to ANSYS Mechanical APDL version 17 (ANSYS, Canonsburg, PA) to reconstruct nodal traction stress with finite element method (FEM) [46]. Traction force was then calculated from the surface traction stress by multiplying with unit mesh area.

### Nucleus Isolation

Nuclei were isolated as previously described [24] based on the methods of [47, 48] with the following modifications: 3T3and N3KO cells were cultured in ultracentrifuge tube inserts coated with 0.1 mg/ml fibronectin for 24 hours before the isolation of nuclei. Cells were incubated with 5 ug/ml Hoechst (diluted in cell culture media) for 30 mins before nucleus isolation. Samples were then centrifuged at 2880 x g for 50 mins in DMEM supplemented with 1% calf serum and 2 ug/ml cytochalasin D at 37 °C using a Beckman-Coulter Optima LE-80 K ultracentrifuge with an SW-28 rotor. Isolated nuclei were collected in inserts coated with 0.1 mg/ml poly-d-lysine positioned beneath the cell-containing inserts. To stabilize the inserts during centrifugation, PDMS was cured in the base of ultracentrifuge tubes to create a flat surface. The volume of isolated nuclei was calculated based on their radii using Image J, assuming a spherical shape.

### Atomic Force Microscopy (AFM)

AFM measurements were made with a Nanowizard 4 (Bruker) mounted on a Leica DMI 6000 B microscope. Before each experiment, a calibration was made by determining the slope of the deflection of the AFM cantilever on a rigid substrate, a plastic petri dish, to determine the deflection sensitivity. Isolated nuclei were compressed using a tipless silicon nitride cantilever (NP-O10, Bruker, Camarillo, CA) with a nominal spring constant of 0.35 N/m at a frequency of 1 Hz and a maximum applied force of approximately 30 nN. Nuclei were distinguished from cells that detached during centrifugation by a near complete overlap between bright field images and Hoechst staining. All measurements were made at room temperature in PBS. 3T3 and N3KO fibroblasts were indented with 1um diameter silica bead attached to silicon nitride cantilever with a spring constant of 0.06 N/m (Novascan, Boone, IA). Measurements were made on 3 points of each cell in the area between the nucleus and the cell periphery at a frequency of 1 Hz and a maximum applied force of 6 nN.

AFM force curves for isolated nuclei were analyzed using the JPK analysis software and fit to a double-contact Hertz model for a sphere being compressed between two planes as previously described [49] while force curves for cells were fit to the Hertz model for a sphere indenting a homogeneous, elastic half-space as described in [50].

### Statistical Analysis

Each experiment was performed at least twice, and each condition was measured for at least 20+ cells unless specifically stated otherwise. Statistical tests for each experiment are stated in the figure caption. Denotations: *, p <= 0.05; **, p <= 0.01; ***, p <0.001; n.s., p >0.05.

## Figures

**SI Fig 1:**
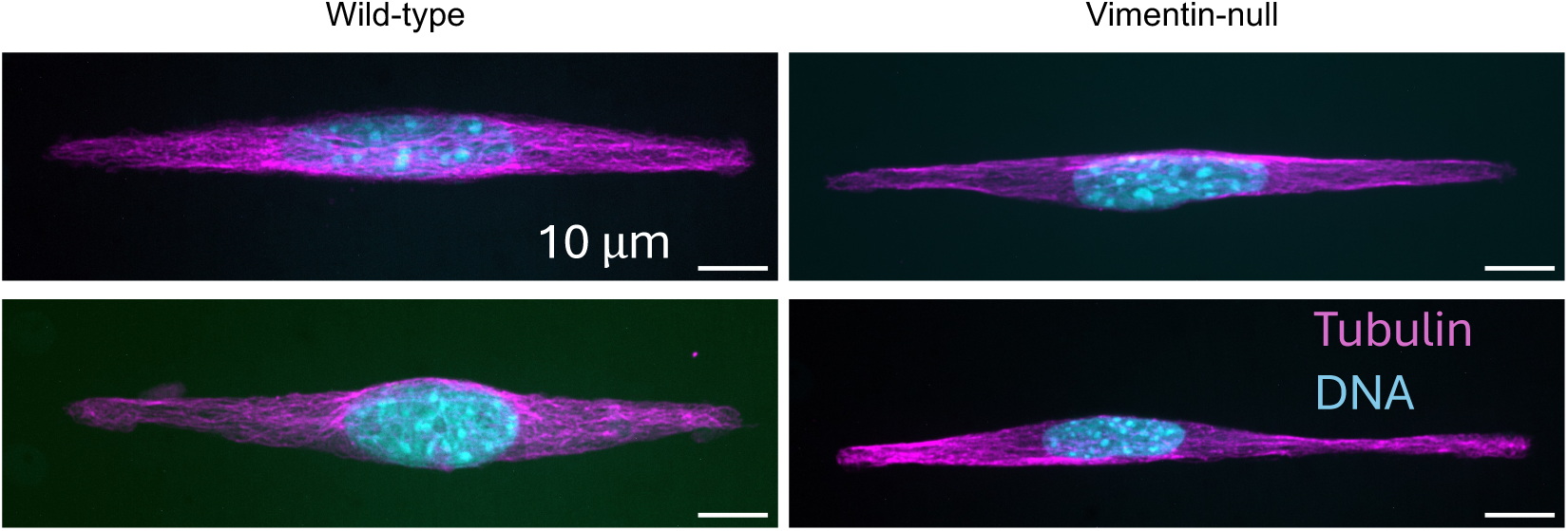
Characteristic images of microtubules on high aspect ratio patterns. Microtubules are colored magenta and DNA is colored cyan.

**SI Fig 2:**
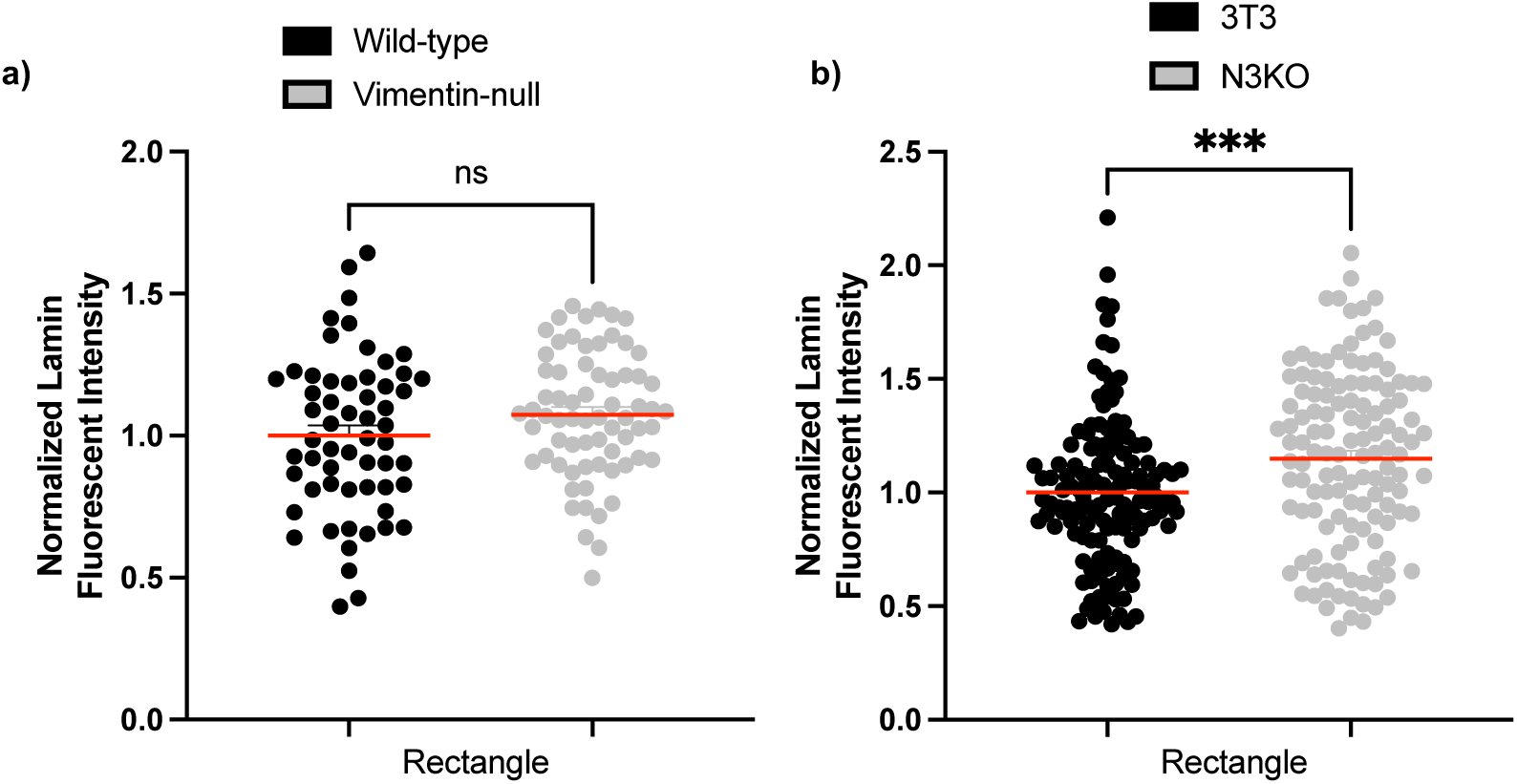
Mean lamin fluorescent intensity for wild-type and vimentin null (a) or 3T3 and N3KO (b) cells on high aspect ratio patterns. Mean intensity is normalized to the wild-type or 3T3 condition for each experiment. Statistical signifigance is tested using an unpaired t-test with Welch’s correction. (For (a) N = 2 experiments with n = 20+ cells per condition; for (b) N=3 experiments with n = 20+ cells per condition).

